# Transcriptional Rewiring of BET Inhibitor Treated Ewing Sarcoma Cells Augments their Dependency on Focal Adhesion Kinase

**DOI:** 10.1101/2025.09.02.673849

**Authors:** Shireen S Ganapathi, Emma D Wrenn, Nicolas M Garcia, Neerja Katiyar, Stephanie I Walter, Aya Miyaki, Jonah Valenti, Yuqi Kang, Marina Chan, Taranjit S Gujral, Elizabeth R Lawlor

## Abstract

Epigenetic dysregulation is a hallmark of cancer. Small molecule inhibitors such as bromodomain and extraterminal (BET) protein inhibitors developed to target epigenetic dependencies have demonstrated significant promise in preclinical models. However, clinical success with epigenetic drugs as single agents has been limited by emergence of tumor cell tolerance and escape, which often occurs due to transcriptional rewiring. Ewing sarcoma (EwS), a bone and soft tissue tumor driven by the EWS::FLI1 fusion oncoprotein, is characterized by profound oncogene-dependent enhancer reprogramming. Thus, epigenetic modifying therapies are of high therapeutic interest. In this study, we sought to elucidate how EwS cells escape BET inhibition to identify biologically informed drug combinations that could be advanced to clinical trials. As expected, EwS cells and xenografts initially responded to BMS-986158, a pharmaceutical grade BET inhibitor, but proliferation was rapidly restored. A kinase inhibitor screen showed that BMS-986158 drug tolerant persister (DTP) cells were sensitive to inhibitors of Focal Adhesion Kinase (FAK), a critical signaling node downstream of extracellular matrix (ECM) engagement. RNA sequencing revealed that DTP cells had been transcriptionally rewired and that mesenchymal signature and ECM remodeling genes were specifically upregulated. Combining BMS-986158 with the FAK inhibitor Defactinib had synergistic effects, reducing EwS cell proliferation, survival, and invasion in vitro, and significantly inhibited tumor outgrowth in vivo. Our studies identify BET and FAK inhibition as a rational combination therapy worthy of further investigation for EwS, and demonstrate that defining emergent mechanisms of epigenetic drug tolerance can identify new vulnerabilities that can be therapeutically targeted.

## Introduction

Epigenetic dysregulation plays a pivotal role in tumor growth, progression, and therapeutic resistance (1). In cancers, this dysregulation often results from mutations in epigenetic regulators, DNA methylation changes, altered histone modifications, and chromatin structural changes (2). Since the introduction and approval of DNA methyltransferase (DNMT) inhibitors (3, 4), significant progress has been made in developing epigenetic-targeting therapies (5). These agents are particularly relevant to pediatric and adolescent malignancies, like Ewing sarcoma (EwS), which exhibit low mutational burden but high epigenetic dysregulation (6, 7). Nevertheless, despite promising preclinical results, the clinical translation of these therapies is hindered by the emergence of drug-tolerant tumor cell populations, particularly in solid tumors.

EwS, a bone and soft tissue malignancy primarily affecting adolescents and young adults, is driven by FET::ETS fusions, most commonly EWSR1::FLI1 (8). EWS::FLI1 broadly reprograms transcriptional networks and acts as a pioneer factor that binds to GGAA microsatellite regions creating *de novo* enhancers and activating genes necessary for oncogenesis (9, 10). As such, there is a high priority to study both the efficacy and escape mechanisms of epigenetic modifying agents in EwS. Of particular relevance are bromodomain and extraterminal (BET) protein inhibitors. These target the BET family of proteins (BRD2, BRD3, BRD4, and BRDT) which function as epigenetic readers and serve as transcriptional activators through binding to acetylated lysine residues (11). Prior preclinical work demonstrated that BET inhibitors have strong anti-tumor efficacy and reverse the EWS::FLI1 transcriptional signature when used as single agents (12–15), as well as and when combined with IGF1R (16) or CDK9 (17) inhibitors. Given this strong preclinical data, EwS patients were included in the first Phase I clinical trial of BET inhibitors in children and adolescents (NCT03936465), which tested the safety of BMS-986158 in patients with relapsed and refractory solid tumors.

A significant challenge to the clinical success of BET inhibitors is the development of drug-tolerant cell states that escape growth-inhibition (18–20). Unlike genetically complex tumors, such as EGFR mutant non-small cell lung cancer (NSCLC), that escape EGFR inhibition by selecting for drug-resistant subclones (21), genomically quiet tumors such as EwS often acquire drug tolerance through non-genomic mechanisms. These include epigenetic and transcriptional rewiring, alternative signaling activation, metabolic reprogramming, and altered interactions with the tumor microenvironment (22, 23). Interrogating these mechanisms is critical for identifying new vulnerabilities that can be leveraged to enhance the effects of epigenetic therapies.

In this study we sought to validate the efficacy of BMS-986158 in preclinical models of EwS and nominate candidate drugs that could be rationally and safely combined with BET inhibition to overcome drug tolerance. Our results confirmed strong anti-tumor efficacy of BMS-986158 as a single agent, with significant cytotoxicity in vitro and tumor growth inhibition in vivo. However, drug tolerant cells rapidly emerged, and these cells adopted mesenchymal morphologies and enhanced invasive potential. Transcriptional profiling showed that BET inhibitor tolerant cells were transcriptionally rewired to upregulate mesenchymal gene signatures, including extracellular matrix (ECM) remodeling genes, that are normally repressed by EWS::FLI1. Independently, a kinase inhibitor screen identified Focal Adhesion Kinase (FAK) as a dependency in BMS-986158 tolerant cells. We therefore tested dual targeting of BET and FAK activity in EwS models and showed that the combination more effectively inhibited tumor growth in vitro and in vivo than either agent alone. Importantly, in contrast to combined BET and PI3K inhibition which was highly toxic, we found the combination of BET and FAK inhibitors to be well-tolerated by mice. Together these studies identify BET and FAK inhibition as a rational combination therapy worthy of further investigation for EwS. Additionally, we demonstrate that defining emergent mechanisms of epigenetic drug tolerance can identify new vulnerabilities that can be therapeutically targeted.

## Methods

### Cell lines

EwS cell lines A673 (RRID:CVCL_0080), CHLA10 (RRID:CVCL_6583), TC32 (RRID:CVCL_7151) were obtained from ATCC and COG cell line repositories. PDX305 is a patient derived xenograft (PDX) line that was derived from a patient with relapsed, widely metastatic disease. PDX305 was generated and profiled as previously described (24) and early passage cells were used for all in vitro experiments. A673 and TC32 were cultured in RPMI 1640 media (Gibco) supplemented with 10% FBS (Atlas Biologicals) and 2mmol/L-glutamine (Life Technologies). CHLA10 and PDX305 were cultured in IMDM media (Fisher) supplemented with 20% FBS, 2mmol/L-glutamine, and 1X Insulin-Transferrin-Selenium-Ethanolamine (Gibco). Cells were cultured at 37°C with 5% CO_2_. Regular *Mycoplasma* and short tandem repeat confirmation were performed every 6 months.

### Generation of BET inhibitor drug tolerant persister cells

2 x10^6^ A673 or CHLA10 cells were plated on 10cm adherent cell culture plates. Cells were incubated continuously with 5nM BMS-986158 for 20 days. Cells were split when confluency reached 70-80% and replated at 2 x10^6^/10mL density. When cell populations stopped proliferating, cell culture medium with fresh drug was replaced every 72hrs. Doubling time of cell populations was calculated using the following equation: time (hrs)*ln2/ln(final cell count/initial cell count). Cells were also collected for subsequent RNA, protein, 3D collagen assays and immunofluorescence assays.

### RT-qPCR

RNA was purified from whole cell lysates using the RNeasy Mini Kit (Qiagen) and cDNA was generated using the iScript cDNA Synthesis Kit (Bio-Rad). qPCR was performed using either Taqman Fast Universal PCR Master Mix (Applied Biosystems) assays or iTaq Universal SYBR-Green Supermix (Bio-Rad) on a Roche Light-Cycler 480 instrument (Roche Applied Science). Average Ct values of three technical replicates were normalized to the geometric mean of two reference genes and relative mRNA expression was calculated by the ddCt method. Fold change was determined by relative mRNA expression of the treatment as compared to control.

### Western blots

Cell pellets were lysed with RIPA buffer (Fisher Scientific) with added protease and phosphatase inhibitors (Sigma). Western blot was performed using the Bio-Rad Mini-PROTEAN Tetra System. Following transfer, nitrocellulose membranes were blocked in Odyssey blocking buffer (LI-COR) for 1 hour. Membranes were washed and incubated rotating overnight at 4°C with primary antibody. Anti-phospho antibodies were additionally incubated for 1 hr at room temperature following overnight incubation. Membranes were then washed three times in TBST for 5 min each and incubated with secondary antibodies (LI-COR IRDye 700CW or 800CW; 1:10,000) for 1 hour at room temperature. Membranes were imaged on a LI-COR Odyssey scanner.

### Dose response assay

Cells were plated in 96-well plates at cell density of 2000-5000 cells/well (depending on each cell lines proliferative rate) in 100uL medium. Serial dilutions of BMS-986158 (supplied directly from Bristol-Meyer Squibb), Copanlisib (Selleckchem Cat # S2802), or Defactinib (VS-6063, Selleckchem, Cat # S7654) were diluted in 100uL media (1:1000 DMSO concentration). Plates were incubated at 37°C with 5% CO_2_ with drug for 72hrs. Cell viability was determined by luminescence using Cell-Titer Glo ATP-based assay (Promega). Luminescence was read using Spectramax iD5 plate reader (1000ms). IC50 were determined using Graphpad Prism (version 10).

### Combination drug synergy assays

Cells were plated in 96-well plates at cell density of 2000-5000 cells/well in 100uL medium. Serial dilutions of BMS-986158 and Copanlisib or BMS-986158 and Defactinib were added in combination at non-fixed ratios, or as single agents, or respective DMSO control to wells. Plates were incubated at 37°C with 5% CO_2_ with drug for 72hrs. Cell viability was determined by the Cell-Titer Glo ATP-based assay (Promega). Luminescence was read using Spectramax iD5 plate reader (1000ms). Total viability of combination treatment or single agents was calculated as treatment viability compared to respective control. Synergy was determined by SynergyFinder Plus (25), specifically calculating Bliss Synergy Scores of Independence (26) and zero interaction potency (ZIP) synergy scores(27). Plots shown in figures are representative of Bliss synergy scores. Supplementary tables have both Bliss and ZIP synergy scores for each individual combination. δ-scores >5 indicate moderate synergy and δ-scores >10 indicates strong synergy. Higher scores indicate stronger confidence that two drugs are synergistic.

### Bulk RNA-sequencing

Poly(A)-capture RNA-seq was performed on A673, TC32, and CHLA10 vehicle (DMSO) cells, A673, TC32, and CHLA10 cells exposed to 5nM BMS-986158 for 72 hrs continuously, and A673 and CHLA10 cells exposed to 5nM BMS-986158 for 20 days continuously (drug tolerant persister (DTP) cells). RNA was purified from whole cell lysates using RNeasy Mini Kit (Qiagen). RNA-seq libraries were prepared by Novogene, and paired end 150 bp sequencing was performed on a Novaseq6000.

### Bulk RNA-seq analysis

Raw sequencing reads were checked for quality using FastQC (RRID:SCR_014583). Adapter sequences were trimmed from the raw sequencing reads using Trim Galore (RRID:SCR_011847). RNA-seq reads were aligned to the GRCh38 assembly using STAR aligner (28) and counted for gene associations against the UCSC genes database. RSeQC (29) was used to check for the quality of the alignments. Differential Expression analysis for RNASeq data was performed using R/Bioconductor package, DESeq2 (30). FDR < 0.05 was used to find transcriptionally regulated genes in response to treatment.

### Pathway analysis

Gene set enrichment analysis (GSEA) was performed for selected gene list sets (31, 32) using the MSigDB (33) database with the Gene Ontology (34, 35) (Biological Process, Cellular Component) and Reactome (36) gene sets using clusterProfiler (37) package. Over-representation analysis (ORA) was additionally performed for the contrasts as detailed in subsequent result sections through Enrichr (38). Resulting enriched genesets and pathways were filtered via a threshold of FDR < 0.05. Dot Plots were created with the enrichplot package in R. Venn Diagrams and barplots were generated using the R packages ggVenndiagram (RRID:SCR_026950) and ggplot2 (RRID:SCR_014601) respectively. GSEA plots were created with the software GSEA_4.3.2.

### Real time proliferation assays

Real-time proliferation was performed on a xCELLigence E-plate 16, on the xCELLigence DP system (Acea Bioscience). Wells were first coated with 0.2% cold gelatin for 10 minutes, which was aspirated off and 5x10^3^ cells/100uL complete media were added. Plates were equilibrated for 1 hr at 37°C in 5% CO_2_ prior to placement in the xCELLigence DP system. Proliferation was measured for up to 96 hours.

### Immunofluorescence

Cells were fixed with 4% PFA in PBS for 10 minutes then washed with PBS. Cells were blocked with 0.2% BSA in PBS for 1 hour at room temperature. Primary antibodies were incubated for 1 hour at room temperature in 0.2% BSA. Alexa-fluor–conjugated secondary antibodies were incubated for 1 hour at room temperature in 0.2% BSA. Signal was compared with matched species-specific IgG controls incubated at the same concentrations. For F-actin staining Alexa Fluor conjugated phalloidin was used (Thermo Fisher, 400X). DAPI was included during secondary antibody incubations to mark nuclei at 2 μg/mL. Coverslips were mounted with ProLong Gold mounting media and allowed to cure overnight at room temperature in the dark. Slides were imaged on a Leica DMi8 Thunder Imager widefield deconvolution microscope.

### Kinase inhibitor (KI) screen

KI screening was performed as described previously (39, 40). The effects of a computationally selected set of 32 KIs on cell growth were assessed using a live-cell imaging assay. Briefly, 5×10^3^ A673 naïve or DTP cell populations were plated per well in 96-well tissue culture plates, allowed to adhere overnight, and then treated with KIs dissolved in DMSO. Drug dilutions were prepared in complete media and applied in triplicate. Cell growth was monitored using the Incucyte imaging system (10× objective), and confluence, defined as the percentage of area covered by cells, was quantified using Incucyte ZOOM 2016B software. The percent confluence at 500 nM, determined from dose-response curves for each inhibitor, was used as the response variable for regression analysis.

### Kinome Regularization (KiR) Modeling

KiR models for both A673 naïve and DTP cell populations were generated as previously described(39, 40). Kinase inhibition profiles of the 32 inhibitors served as explanatory variables, and corresponding cell response data were used as response variables in elastic net–regularized multiple linear regression models. Models were built using custom R scripts (https://github.com/FredHutch/KiRNet-Public) with the glmnet package. Leave-one-out cross-validation (LOOCV) was used to select the optimal penalty factor (λ), and models were computed across 11 values of α (ranging from 0 to 1) to balance Least Absolute Shrinkage and Selection Operator (LASSO) and Ridge regularization. Model performance was evaluated using LOOCV error and root-mean-squared error (RMSE) of predictions.

### In vivo experiments

All experimental procedures adhered to the *Guide for the Care and Use of Laboratory Animals* and were approved by Seattle Children’s Research Institute IACUC. The institute is fully AAALAC-accredited and has a Public Health Service approved Animal Welfare Assurance. BMS-986158 (supplied directly from Bristol Meyer Squibbs (BMS)) was diluted in the following vehicle as per guidance from BMS: polyethylene glycol 400 (PEG 400),Propylene glycol (PG), d-alpha tocopheryl polyethylene glycol 1000 succinate (TPGS), Ethanol (EtOH): 55/25/15/5 w/w/w/w PEG400/PG/TPGS/EtOH to a concentration of 3.0mg/mL. Mice were dosed 1.6mg/kg/dose via oral gavage (OG) twice a day (BID) 5 days on/2 days off. Copanlisib (MedChemExpress, Cat # HY-15346), was dissolved in 0.05% Trifluoroacetic acid (TFA) and NaCl to a concentration of 2mg/mL. Mice were dosed 14mg/kg/dose IV every other day, 3x/week. Dose was reduced to 10mg/kg/dose IV every other day, 3x/week due to toxicity. Defactinib (Selleckchem, Cat # S7654) was diluted in 5% DMSO and 55/25/16/5 w/w/w/w PEG400/PG/TPGS/EtOH to a concentration of 20mg/mL and mice were dosed 50mg/kg/dose via OG BID 5 days on 2/days off. Vehicle mice received OG of diluent 55/25/16/5 w/w/w/w PEG400/PG/TPGS/EtOH at same dosing schedule and route as experimental groups. For all mice experiments, equal numbers of male and female mice were used. For subcutaneous experiments 1x10^6^ cells were injected with 1:1 Matrigel (Corning) into the flank of the mice. When tumors reached 100m^3^, mice were randomized to a treatment arm. Body weight and tumor volume, by caliper measurements (V=(L x W^2^)/2), were recorded 3 times per week. Mice were sacrificed when tumor diameter reached 2 cm in longest dimension, or sooner if >15% weight loss or additional humane endpoints were reached. For tail vein experiments, 1x10^6^ GFP/Luciferase+ expressing cells were injected via tail vein in 100uL PBS. In Vivo Imaging System (IVIS) was performed at 1hr to confirm successful injection. Mice started treatment 72hrs after injection. Treatment was continued for up to 8 weeks. Serial bioluminescence signal was monitored weekly via IVIS. Engraftment was defined as a luminescence signal greater than 1 x 10^7^ on two consecutive measurements.

### 3D rat tail collagen invasion assays

50,000 viable cells/mL in 4mL complete media were plated in ultra-low-attachment 6 well plates overnight (Thermo Fisher Scientific 07-200-601) to form spheroids. Debris and single cells were removed by centrifuging samples for 10 seconds at 300 g the following day. Rat tail collagen (Gibco) was prepared at 1.8 to 2 mg/mL in 0.08 N acetic acid, supplemented with 10X DMEM (final concentration 1X), neutralized with 1 mol/L NaOH and allowed to polymerize on ice for 45 to 60 minutes until fibers formed (41). Spheroids and collagen gels were mixed and plated as 80 μL droplets and allowed to polymerize for 30 minutes at 37°C, at which point 1 mL of complete IMDM media (supplemented with 20% FBS, 1X L-Glutamine, 1X ITS-X) was added. 20 μL of polymerized rat tail collagen was first applied to the bottom of a tissue culture–treated 24wp plate to prevent spheroids in the overlying collagen dome from attaching to or spreading along the plate surface. Spheroids in rat tail collagen were cultured overnight. The following morning spheroids were treated with vehicle, 5nM BMS-986158, 5μM Defactinib, or the combination. Spheroids were cultured in treatment for 6 days and then assessed by phase contrast microscopy and fixed for 15 minutes with 4% PFA in PBS. Invasion was defined as protrusive cells or multicellular strands extending from the spheroid border. Cell viability in a 3D matrix was subsequently assessed after 6 days, with the 3D CellTiter-glo 3D Cell Viability Assay (Promega). Equal volume of CellTiter-Glo was added to media in each well and put on rotator for 40 min at room temperature to lyse spheroids and homogenize gels. Well contents were transferred to an opaque white-walled 96 well plate and luminescence was read using Spectramax iD5 plate reader (1000ms). Total percent viability was normalized to control.

### Statistics and software

GraphPad Prism (RRID:SCR_002798) version 10 was used for statistical analysis. Kaplan-Meier Curves were analyzed using Log-rank (Mantel-Cox) test. Student’s t-test was used to determine significant differences between two groups and one-way ANOVA with Tukey’s Multiple Comparison for any group analysis greater than two unless specified otherwise. All error bars are representative of standard error of mean of at least 3 biological replicates. Statistical significance: ns, p>0.05; * p<0.05, ** p<0.01, *** p<0.001, **** p<0.0001

### Data and code availability

Sequencing data generated by this study will be available in the NCBI Gene Expression Omnibus database upon publication. Analyses in R will be available upon publication at github.com/LawlorLab.

## Results

### Clinical grade BET inhibitor BMS-986158 slows proliferation of EwS in vitro and in vivo and inhibits EWS::FLI1-dependent gene activation

To first determine if the clinical grade BET inhibitor BMS-986158 inhibits EwS tumor growth we tested the drug as a single agent in in vitro and in vivo models. We exposed both established and PDX-derived EwS cell lines to increasing doses of BMS-986158 and assessed viability following 72 hrs of drug exposure. As shown, IC50s for the drug across models were reproducibly in the low nanomolar range (**Fig 1A**). To more specifically evaluate the effects of BMS-986158 on cell proliferation, A673 and CHLA10 cells were exposed to ∼IC50 and ∼IC75 doses of drug, and cell expansion was monitored using real time proliferation assays (as described in Methods). Proliferation was significantly and reproducibly reduced at both concentrations (**Fig 1B**). Next, we tested whether BMS-986158 would alter the growth of established EwS tumor xenografts. Growth of both A673 and TC32 subcutaneous tumors was delayed by BMS-986158 and this resulted in prolonged survival of the mice. TC32 xenograft average survival post treatment was 18 days for vehicle and 30.5 days for treatment group (p=0.0012) (**Fig 1C**). A673 xenograft average survival was 20 days post treatment for vehicle and 27 days for treatment (p=0.0174) (**Supplementary Fig S1A**). While delayed tumor growth was observed in both xenograft models (**Supplementary Fig S1B, C**), treatment groups eventually reached tumor endpoint (**Supplementary Fig S1D,E**), highlighting that BMS-986158 as a single agent slows growth but is not sufficient for tumor control. BMS-986158 was well tolerated, and no significant weight loss was observed among treated mice (**Supplementary Fig S1F**). Additionally, at necropsy gross examination revealed no evidence of end organ toxicity.

**Figure 1.**
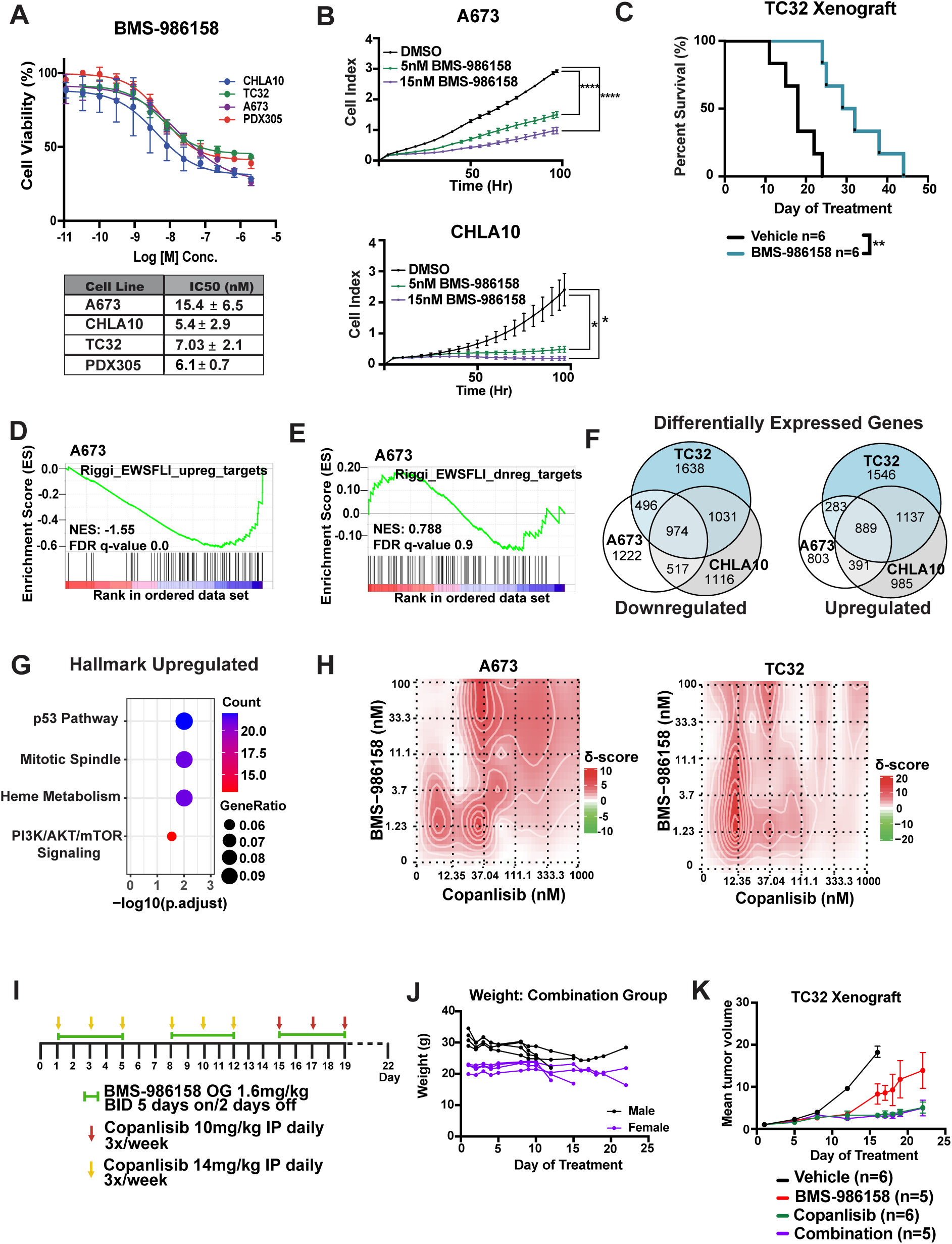
BMS-986158 slows proliferation of EwS in vitro and in vivo and inhibits EWS::FLI1-dependent gene activation. **A.** Dose response curves of multiple EwS cell and PDX lines. Cells were treated with serial dilutions from 2uM to 0.03nM, or relevant DMSO controls. Y-axis represents % viability as compared to DMSO control. Table below shows IC50 values (n=3-4 individual experiments with three technical replicates per condition). **B.** xCELLigence Real Time Cell Proliferation Assays (n=2, with 3 technical replicates per experimental condition). **C.** Kaplan-Meier overall survival curves of TC32 xenografts treated with BMS-986158 or vehicle (p=0.0012, log-rank test). RNA-seq of A673 cells treated with 5nM BMS-986158 for 72hrs as compared to DMSO control. GSEA of 100 enhancer-driven genes **D.** activated or **E.** repressed by EWS::FLI1(10). **F.** Venn diagram of statistically significant (padj<0.05) differentially expressed genes that are, left-downregulated (Log2Fold<0) or right-upregulated (Log2Fold>0) with 72hrs treatment of 5nM BMS-986158 as compared to vehicle. **G.** Hallmark (Enrichr) over-representation analysis of the 889 genes that are upregulated in A673, CHLA10, and TC32 following 72hr treatment with BMS-986158. **H.** Bliss Synergy(26) plots of BMS-986158 (0.41nM-100nM, serial dilutions 1:3) and Copanlisib (12.35nM-1uM, serial dilutions 1:3) exposed A673 and TC32 cells (n=3, with three technical replicates per condition). TC32 subcutaneous xenograft in vivo experiment **I.** Schematic of experiment with dosing, route, and schedule of all individual drugs. Experiment was stopped at Day 22 given toxicity. **J.** Weight of combination group over time. **K.** Mean tumor volume excluding mice in the combination group who were humanely euthanized at D14 given weight loss >15%.

It has previously been established that the anti-tumor effects of BET inhibitors in EwS are largely mediated by the effects of BRD4 inhibition on EWS::FLI1-dependent oncogenic gene signatures (12, 13, 15). Specifically, exposure of EwS cells to the tool compound JQ1 inhibited both EWS::FLI1-mediated gene activation and also partially reversed fusion-dependent gene repression (12, 13, 15). To investigate the effects of BMS-986158 on the EWS::FLI1 transcriptional signature we performed bulk RNA-seq on A673, CHLA10, and TC32 cells that had been exposed to vehicle or 5 nM BMS-986158 (∼IC50) for 72 hrs (**Supplementary Table 1**). In keeping with published data, exposure of EwS cells to BMS-986158 resulted in strong reversal of the EWS::FLI1-activated signature (**Fig 1D, Supplementary Fig S1G**). In contrast to JQ-1 however, short term exposure of EwS cells to BMS-986158 did not induce broad activation of EWS::FLI1-repressed genes (**Fig 1E, Supplementary Fig S1H**). Thus, the clinical grade BET inhibitor BMS-986158 inhibits the proliferation of EwS tumor cells both in vitro and in vivo and this is associated with strong inhibition of EWS::FLI1-dependent gene activation.

### BET and PI3K inhibitors are synergistic but induce systemic toxicities in vivo when used in combination

Consistent with prior reports (16, 17, 20), our studies demonstrate that BET inhibitors will not be effective therapies for EwS tumors if used as single agents. To identify drugs that could augment the efficacy of BMS-986158, we sought to define the molecular features of EwS tumor cells that escaped the anti-proliferative effects of BET inhibition. To address this, we interrogated the transcriptional profiles of cells that remained viable and that continued to proliferate, albeit much more slowly, after 72 hours exposure to BMS-986158.

As anticipated, given its broad effects on gene regulation, BMS-986158 had a marked impact on the transcriptomes of all three EwS cell lines beyond its effects on the EWS::FLI1 activation signature. Highlighting the documented intertumoral heterogeneity of EwS models (24), acute exposure to BMS-968158 had distinct effects on each of the cell lines. Nevertheless, a subset of genes was commonly altered across all three models compared to their respective untreated controls (**Fig 1F**). Over-representation analysis identified enrichment of the PI3K/AKT/mTOR pathway among genes that were reproducibly upregulated in cells that survived acute BET inhibition (**Fig 1G**). Other top enriched pathways among upregulated genes included Rho GTPase signaling and cellular processes involved in trafficking and post translational modification (**Supplementary Fig S2A-B**).

Enrichment of the PI3K/AKT/mTOR pathway among upregulated genes was notable given prior studies of EwS (16) and neuroblastoma (42) that also identified IGF1-R and PI3K pathways as vulnerabilities in BET inhibitor exposed cells. Therefore, we tested BMS-986158 in combination with the clinical grade PI3K inhibitor Copanlisib. As a single agent, Copanlisib was highly effective at inhibiting viability of A673, TC32, and CHLA10 cells in vitro **(Supplementary Fig S2C)**. In addition, the potency of both Copanlisib and BMS-986158 was enhanced when they were applied in combination, and strong synergy was observed across all three models at sub-IC50 doses of each drug (**Fig 1H, Supplementary Fig S2D**). Synergy plots are representative of Bliss synergy (26) modeling scores, ZIP synergy scores (27) were also calculated (**Supplementary Table 2**).

We next evaluated the combination in vivo. Subcutaneous TC32 xenografts were established in NSG mice, and mice were randomized to receive vehicle, BMS-986158, Copanlisib, or the combination once tumors reached 100mm^3^ (**Fig 1I**). Although each single agent was well tolerated, greater than half of the mice in the combination treatment group (5/9) exhibited weight loss after two cycles, with three out of the nine mice experiencing weight loss >15% from baseline requiring humane euthanasia (**Fig 1J**). Treatment was held for the surviving mice in the combination group until their weight returned to baseline. Dosing was resumed with a 25% reduction in Copanlisib. Despite reduced dosing, ongoing toxicity led to humane euthanasia of all remaining combination mice by Day 22 of treatment. Thus, although a reduction in average tumor volume (**Fig 1K**) and prolonged survival (**Supplementary Fig S2E**) were observed in mice treated with both BMS-986158 and Copanlisib, this combination proved too toxic for further evaluation in NSG models. Given this result, and emerging data around dose-limiting hematologic toxicities of Copanlisib in heavily pre-treated patients (43), we sought to identify other agents that could augment the anti-tumor efficacy of BMS-986158 without creating additional toxicity.

### BMS-986158 treated EwS cells reestablish growth potential despite continued inhibition of EWS::FLI1-dependent gene activation

As shown in Figure 1, exposure of EwS cells to single agent BMS-986158 for 72 hours was profoundly cytostatic, though viable cells remained at the end of drug treatment. Slowed growth was also initially observed in BMS-986158 treated tumors in vivo. However, after two weeks, subcutaneous tumors appeared to escape the anti-tumor effects of BMS-986158 and proceeded to grow rapidly to endpoint. Using slope as a surrogate for tumor velocity, we performed linear regression of the change in average tumor volume over time to quantify changes in growth rate. As shown, during the first 15 days of treatment, the rate of growth of BMS-986158 treated A673 and TC32 tumors was significantly slower than their respective controls (**Fig 2A**). In contrast, between Day 18 and tumor endpoint the rate of growth was statistically equivalent between treated and untreated tumors (**Fig 2B-C**). We therefore tested if the inhibitory effects of BMS-986158 on EwS cells growth in vitro would be similarly lost following prolonged drug exposure. Consistent with in vivo studies, after a two- and half-week period of profound growth inhibition, BET inhibitor treated EwS cells resumed proliferating at the same rate as vehicle treated cells by 20 days (**Fig 2D**).

**Figure 2.**
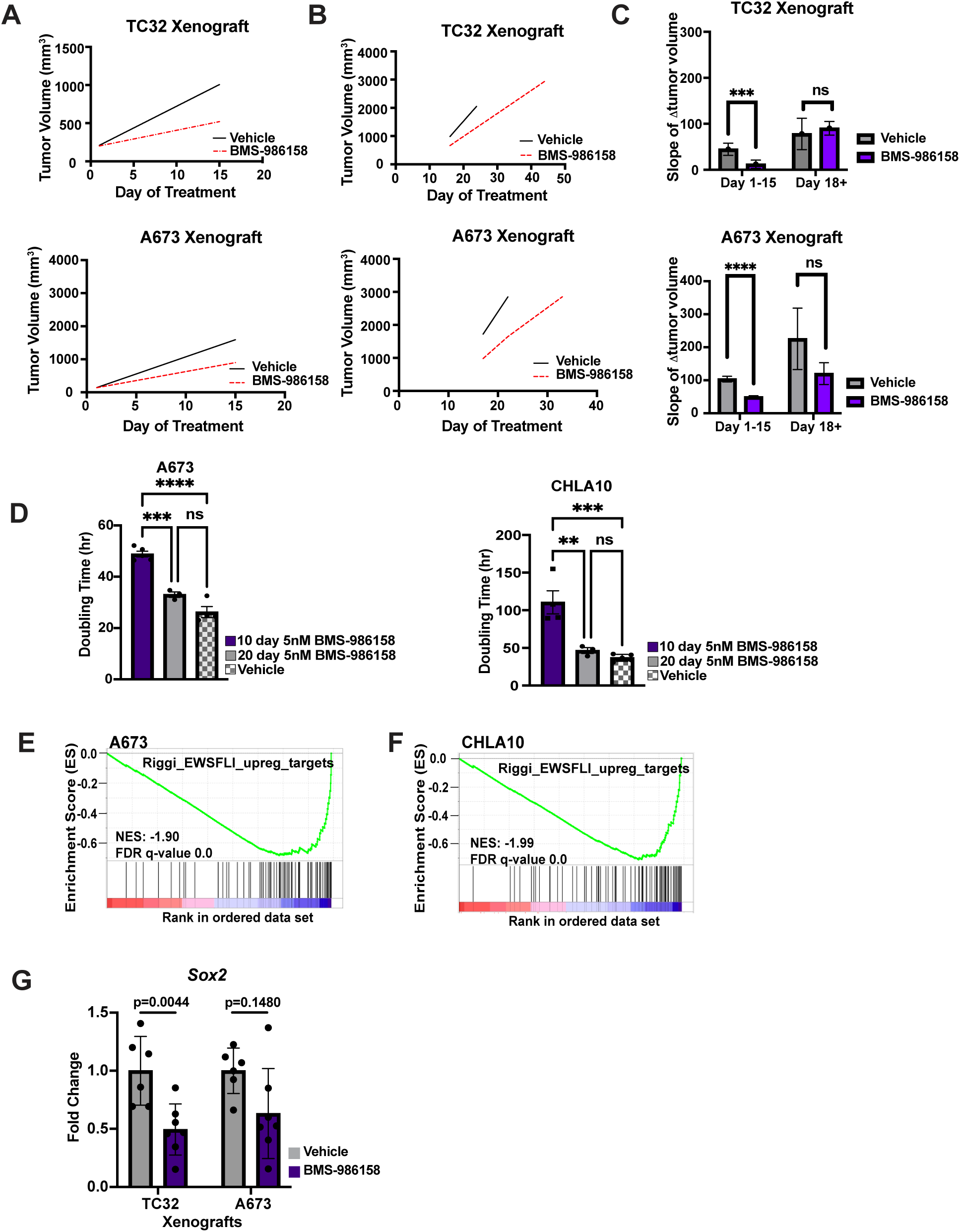
BMS-986158 treated EwS cells reestablish growth potential despite continued inhibition of EWS::FLI1-dependent gene activation. Average tumor volume of TC32 and A673 subcutaneous xenografts treated with BMS-986158 or vehicle from **A.** Day 1-15, and **B.** after 18 days of treatment until tumors reached endpoint. Linear regression of tumor growth velocity was performed and the slope was calculated as a surrogate for tumor velocity. Plotted for BMS-986158 or vehicle for days 1-15 and after day 18 for **C.** TC32 and A673 xenografts. **D.** Doubling time of A673 and CHLA10 cells exposed continuously in culture to 5nM BMS-986158. RNA-seq of A673 and CHLA10 cells treated with 5nM BMS-986158 for 20 days as compared to DMSO control. GSEA of 100 enhancer-driven genes **E.** activated or **F.** repressed by EWS::FLI1 (10). G. *Sox2* gene expression by RT-qPCR from A673 and TC32 subcutaneous xenograft tumors extracted at tumor endpoint.

Having demonstrated that EwS tumors and cells reestablish baseline levels of proliferation after prolonged exposure to BMS-986158, we hypothesized that growth-restored EwS cells had escaped the effects of BET inhibition and had reactivated the EWS::FLI1-dependent oncogenic gene signature. To test this, we used RNA-seq to evaluate the transcriptomic profiles of proliferating EwS cells that had been exposed to 5nM BMS-986158 continuously for 20 days. In contrast to our prediction, the EWS::FLI1-activated gene signature was still significantly repressed in these proliferating cells (**Fig 2E**). In addition, expression of the direct EWS::FLI1 target gene *SOX2* was reduced in EwS xenografts that had escaped the growth inhibitory effects of BET inhibition (**Fig 2F**). Thus, these studies together reveal that EwS cells that survive the acute growth inhibitory effects of BET inhibition, can rapidly restore their proliferative potential in vitro and in vivo despite continued BET inhibitor-mediated suppression of the EWS::FLI1 activated gene signature.

### Kinase inhibitor screen identifies Focal Adhesion Kinase (FAK) as a dependency in BET inhibitor DTP cells

Cancer cells that undergo phenotypic changes in the setting of continued drug exposure and suppression of oncogenic signatures have been termed drug tolerant persister (DTP) cells (44). Therapeutic targeting of DTP cells has the potential to augment the efficacy of single agents and improve patient outcomes (23). Given that EwS cells are initially profoundly sensitive to the effects of BET inhibition, we posited that illumination of specific vulnerabilities in BET inhibitor DTP cells would identify drugs that could be rationally combined with BMS-986158 to enhance and prolong its tumor inhibitory effects. We elected to test this hypothesis using two orthogonal methods: (i) via a drug screening approach and (ii) by interrogating the transcriptional profiles of emergent DTP cell populations.

First, we performed a polypharmacology-based kinome screen to identify kinases that may be distinct vulnerabilities in DTP cell populations (**Fig 3A**). We implemented Kinase inhibitor Regularization (KiR), an approach that takes advantage of the fact that most kinase inhibitors (KIs) are not highly specific and affect multiple targets. Our curated set of 32 KIs collectively covers over 90% of the kinome, enabling broad functional analysis. Using machine learning to integrate drug response data, KiR identifies key kinases driving a phenotype, accounting for redundancy and compensatory pathways often missed by single inhibitors. This method has been extensively validated and applied across various biological systems, including cancer cell migration (45–47), cancer cell growth (39, 40), COVID-19 (48), malaria (49), and cytokine signaling (48, 50). We performed this screen on both A673 naïve and DTP cell populations. Each population was exposed to DMSO or each of the 32 computationally selected kinase inhibitors at four concentrations. Cell survival was calculated by measuring area per image using Incucyte Imaging System after 96 hrs. Initial results demonstrated sensitivity of naïve and DTP cell populations to many clinical grade compounds; however, DTP cell populations were generally more resistant to kinase inhibitors than naïve cells. To improve the accuracy of the model, an additional 8 compounds not included in the upfront screen were tested in naïve and DTP cells. These 8 compounds were either (i) predicted to have a strong effect, (ii) predicted to have a moderate effect, or (iii) predicted to not have an effect (**Supplementary Table 3**). Subsequently, a predicted set of top key kinases in naïve, DTP cells, and shared amongst both were identified. In particular, three kinases were identified by KiR as being dependencies in DTP cells: Focal Adhesion Kinase (FAK), Misshapen-like kinase 1 (MINK1), and Dual-specificity tyrosine phosphorylation-regulated kinase 1 (DYRK1) (**Fig 3B**). KiR identified the established EwS therapeutic targets, IGF1-R (16) and CDK4/6 (51), as shared dependencies in both naïve and DTP cell populations, revealing the critical importance of these pathways in EwS cells that are otherwise predicted to be highly resistant to kinase inhibition. Further we performed KirNet (52), a network-based model that integrates protein-protein interactions from established databases, with results from KiR to predict the critical interaction of both kinase and non-kinase proteins to the phenotype observed. Importantly, KirNet Modeling further nominated FAK as a dependency in DTP cell populations (**Fig 3C**). FAK signaling mediates tumor cell survival, adhesion/migration, and proliferation in multiple cancers, and FAK inhibition was previously shown to slow the growth of EwS tumor xenografts (53). Thus, integration of this polypharmacology-based kinome screen and computational network modeling, identified FAK inhibition as a compelling strategy to pursue in combination with BMS-986158.

**Figure 3.**
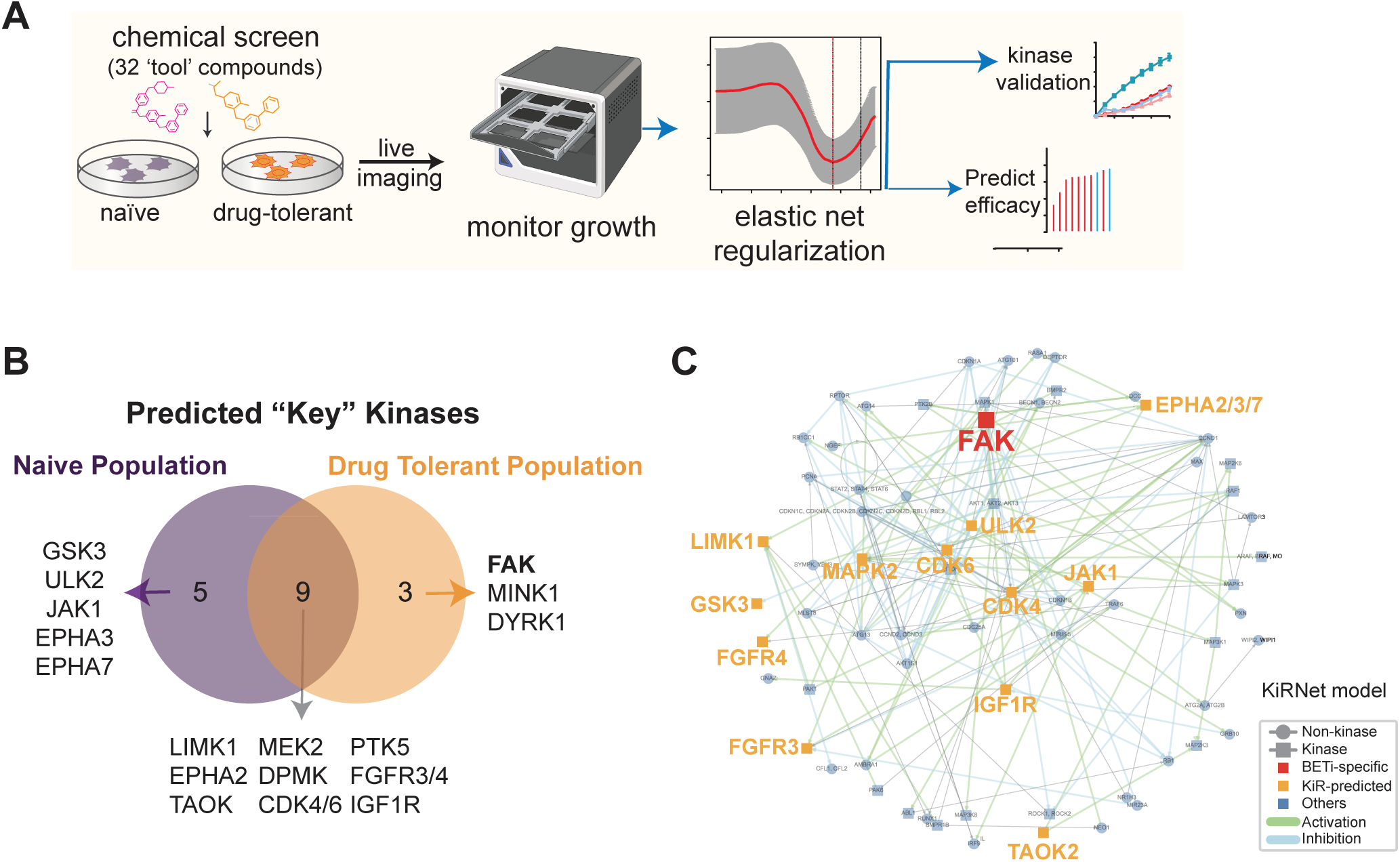
Kinase inhibitor screen identifies Focal Adhesion Kinase (FAK) as a dependency in BET inhibitor DTP cells. **A.** A673 naïve and DTP cell populations were treated with a panel of 32 computationally selected kinase inhibitors and confluence monitored by live imaging. Drug response data were used to train elastic net regression models, which predicted kinases driving cell survival. **B.** Venn diagram showing predicted “key” kinases in naïve (purple) and drug-tolerant (orange) populations. Nine kinases were shared between the two states, while FAK (PTK2) was uniquely predicted in the DTP population. **C.** Network map of predicted kinase regulators from KiRNet modeling. Nodes represent KiR-predicted kinases (orange) or non-kinase interactors (grey circles), and edges indicate known regulatory interactions (green: activation; blue: inhibition). FAK is highlighted as a central node in the DTP-specific network, suggesting a critical role in drug tolerance.

### BET inhibitor-tolerant Ewing sarcoma cells are transcriptionally and phenotypically rewired

As a second and orthogonal approach to identifying therapeutic vulnerabilities in BET inhibitor DTP cells, we interrogated the transcriptional profiles of BET inhibitor treated EwS cells over time. Studies in acute myeloid leukemia (AML) (19), NB (42), and breast cancer (20) showed that DTP cells escape the cytostatic effects of BET inhibition by undergoing epigenetic and transcriptional rewiring to new transcriptional states. We observed that like acute growth arrested cells, DTP cells demonstrate significant transcriptomic alterations as compared to naïve populations (**Fig 4A**). Given the striking difference in proliferation that was evident between acutely treated and DTP cells, we focused on investigating changes in gene expression profiles that had emerged as cells moved from growth arrested to proliferative states. As shown, in both A673 and CHLA10 cells, the expression of hundreds of genes was altered in DTP cells compared to acutely treated cells, supporting the hypothesis that the proliferative potential had been restored via transcriptional rewiring (**Fig 4B**).

**Figure 4.**
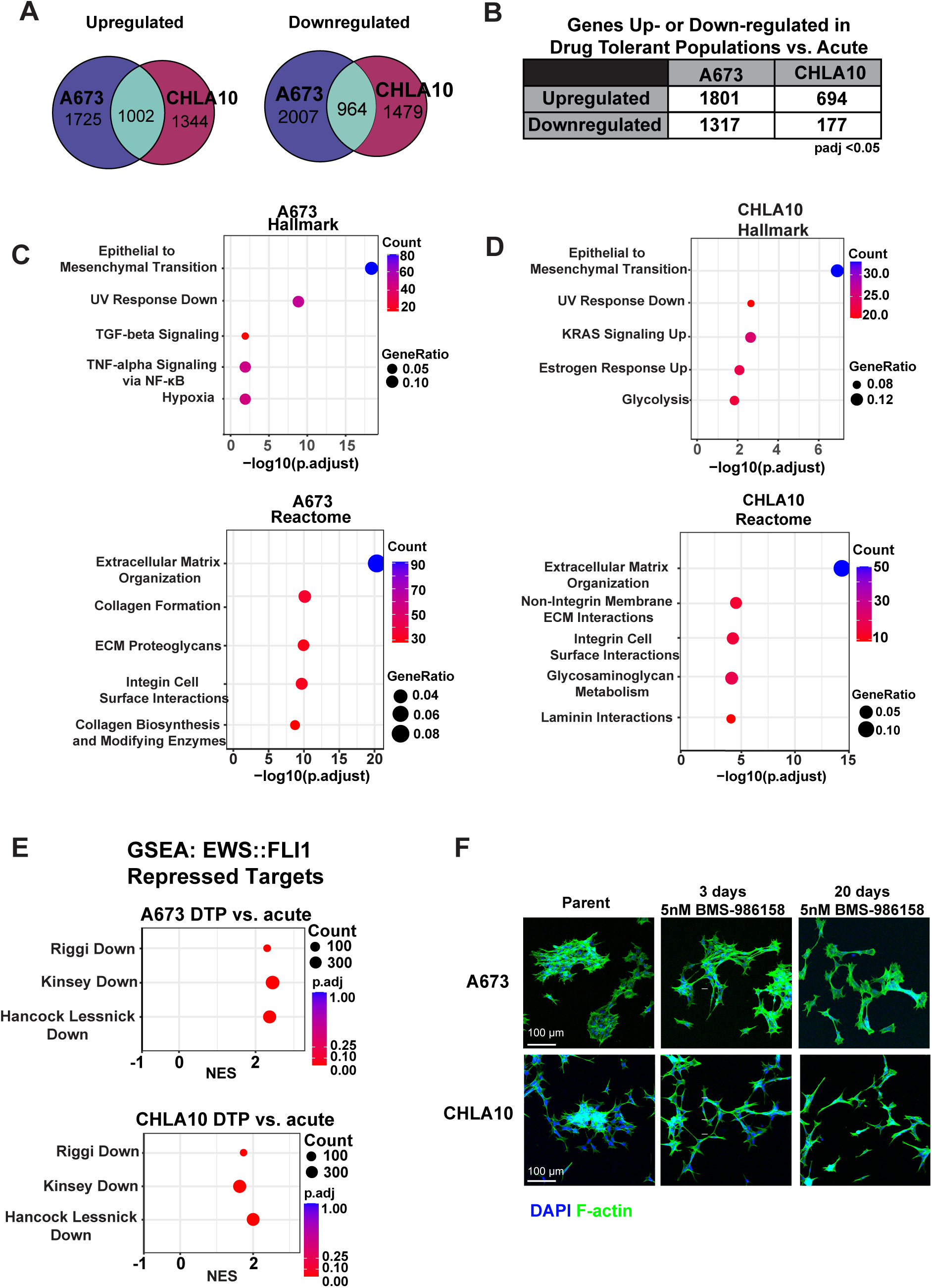
BET inhibitor-tolerant Ewing sarcoma cells are transcriptionally and phenotypically rewired. **A.** Venn diagram of statistically significant (padj<0.05) differentially expressed genes that are, left-upregulated (Log2Fold>0) or right-downregulated (Log2Fold<0) with 20 days of treatment with 5nM BMS-986158 as compared to vehicle. Differential expression by RNA-seq of cell lines treated with 5nM BMS-986158 for 20 days as compared to 5nM BMS-986158 for 72hrs was performed. **B.** Table of statistically significant (padj<0.05) differentially expressed genes that are either upregulated (Log2Fold>0) or downregulated (Log2Fold<0). **C.** Hallmark (Enrichr) and **D.** Reactome (Enrichr) over-representation analysis of the 1801 genes in A673 cells and 694 genes in CHLA10 cells that are significantly upregulated. **E.** GSEA of the repressed EWS::FLI1 signatures (10, 66, 67). **F.** DAPI and F-actin staining of A673 and CHLA10 cells treated with DMSO, 72hrs or 20 days of 5nM BMS-986158 (representative images of n=2).

We performed over-representation analysis on the group of genes that were statistically significantly upregulated in the DTP cells. Strikingly, the most enriched gene signatures in both CHLA10 and A673 DTP cells was similar, suggesting that despite their transcriptomic heterogeneity, the two cell lines may have employed a common mechanism of BET inhibitor escape. Among genes that were reproducibly upregulated in DTP cells, epithelial to mesenchymal transition (EMT) and extracellular matrix (ECM) organization (**Fig 4C-D**) genes were overrepresented suggesting that DTP cells had been transcriptionally rewired to more mesenchymal cell states. Gene Ontology analysis of differentially upregulated genes also supported this conclusion (**Supplementary Fig S3A-B**).

EwS cells exist on a transcriptional continuum between neuroepithelial and mesenchymal states that is in large part determined by the expression level and transcriptional activity of the EWS::FLI1 fusion (54). Significantly, many of the ECM pathway genes that were reproducibly upregulated in BET inhibitor DTP cells are normally repressed by EWS::FLI1. Thus, we used supervised analysis to assess the impact of BET inhibition on the EWS::FLI1-repressed gene signature over time. Despite having little impact in the short term (72 hrs treatment), BET inhibitor DTP cells (20 days treatment) showed a highly significant and reproducible upregulation of EWS::FLI1-repressed genes relative to acutely treated, growth arrested cells (**Fig 4E**). Phalloidin staining confirmed progressive alteration of BMS-986158 treated cells to more mesenchymal phenotypes and at 20 days DTP cells were exclusively mesenchymal with respect to cell shape, reduction in cell:cell contacts, and dominant F-actin rich cell protrusions (**Fig 4F**). Progressive loss of EWS::FLI1-dependent gene repression in DTP cells was not accompanied by progressive changes to EWS::FLI1-mediated gene activation (**Supplementary Fig S3C**) or altered expression of the fusion total transcript (**Supplementary Fig S3D)** or protein (**Supplementary Fig S3E)**.

Together these data show that emergence of BET inhibitor DTP EwS cells is accompanied by profound transcriptional rewiring and this reproducibly induces upregulation of mesenchymal and matrisomal genes that are normally repressed by EWS::FLI1. Engagement of matrisomal proteins by integrins on cytoplasmic membranes of tumor cells induces activation of FAK(55). Thus, results from two unbiased approaches, a kinome inhibitor screen and transcriptional profiling of BET inhibitor DTP cells, converged on FAK as a potential vulnerability in EwS cells that escape the growth inhibitory effects of BET inhibition.

### The combination of BET and FAK inhibition is synergistic in vitro

To begin to test if BMS-986158-tolerant cells would be susceptible to FAK inhibition, we first assessed levels of FAK activation in naïve and BMS-986158 treated cells over time. As shown, phosphorylation of FAK at its autophosphorylation site, Y397, and at the Src substrate Y925, increased over time in both A673 and CHLA10 cells and was most pronounced in proliferating DTP cells (**Fig 5A**). To determine if FAK activation was required for proliferation of DTP cells, we exposed cells to the clinical grade FAK inhibitor, Defactinib. Proliferation of both treatment naïve and DTP cells was inhibited by Defactinib in a dose dependent manner (**Fig 5B**). In DTP cells this was accompanied by a return of FAK phosphorylation to baseline levels (**Fig 5C**).

**Figure 5.**
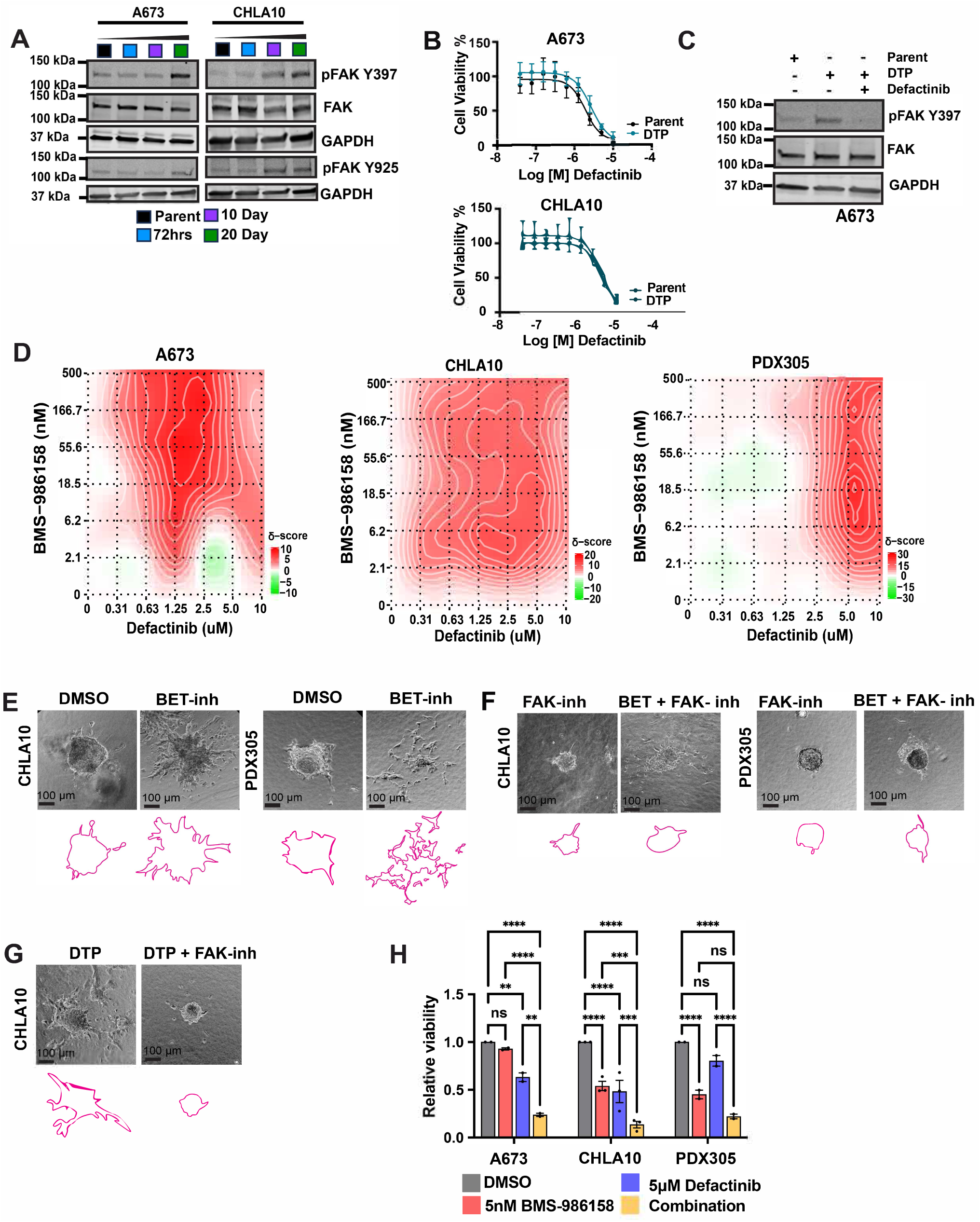
The combination of BET and FAK inhibition is synergistic in vitro. **A.** Immunoblot of pFAK Y397, pFAK Y925 and total FAK with GAPDH used as loading control in A673 and CHLA10 cells treated with 72hrs, 10 days, or 20 days of 5nM BMS-986158. **B.** Dose response curves of A673 and CHLA10 naïve and DTP cells. Cells were treated with serial dilutions from 20nM to10uM or relevant DMSO controls. **C.** Immunoblot of pFAK Y397 and total FAK of A673 naïve, DTP, and DTP cells treated with 2.5uM Defactinib. **D.** Bliss Synergy (26) plots of BMS-986158 (2.06nM-500nM, serial dilutions 1:3) and Defactinib (0.312uM-10uM, serial dilutions 1:3) exposed to 72 hrs of combination, single agent, or DMSO in A673, CHLA10, and PDX305 cells. Invasion through a collagen matrix of CHLA10 and PDX305 spheroids exposed to **E.** 5nM BMS-986158 or DMSO and **F.** 5uM Defactinib or combination of 5nM BMS-986158 and 5uM Defactinib. **G.** CHLA10 drug tolerant spheroids were embedded in rat tail collagen with or without 5uM Defactinib. **H.** A673, CHLA10, and PDX305 spheroids plated on a rat tail collagen matrix were treated with DMSO, 5nM BMS-986158, 5nM Defactinib, or the combination. Following 6 days of treatment, cell viability was assessed with CellTiter Glo (n=2-3, error bars =SEM).

Next, we tested whether combining Defactinib with BMS-986158 would enhance the therapeutic efficacy of the BET inhibitor. We exposed A673 and CHLA10 EwS cell lines to increasing doses of Defactinib, BMS-986158, or the two drugs in combination, and measured viability. In addition, we similarly tested an early passage PDX-derived cell line that was established from a patient with highly aggressive relapsed and drug refractory disease (24). Synergy of the combination was determined using both Bliss Synergy (26) and ZIP (27) modeling (**Supplementary Table 4**). At multiple doses, including IC50 or sub IC50 doses of each individual drug, the combination of Defactinib and BMS-986158 had strongly synergistic effects resulting in significant inhibition of cell viability (**Fig 5D**). We then tested if the drug combination would affect the mesenchymal phenotypes that we observed in BET inhibitor DTP cells. EwS tumor cell spheroids were established in 3D collagen matrices and the effects of drug exposure on cell invasion were measured. Consistent with their transition to more mesenchymal cell states, BMS-986158 treated cells demonstrated an enhanced capacity to invade compared to drug naïve populations (**Fig 5E, Supplementary Fig S4A**) and this early invasive phenotype was blocked by Defactinib (**Fig 5F, Supplementary Fig S4B)**. Importantly, spheroids derived from DTP cells that were resistant to BMS-986158-dependent growth inhibition were highly invasive in 3D collagen, and their invasive capacity was also blocked by Defactinib (**Fig 5G, Supplementary Fig S4C**). Viability of 3D spheroids was also more profoundly inhibited by BMS-986158 and Defactinib in combination compared to either drug alone (**Fig 5H**). In summary, these in vitro studies confirmed our prediction that inhibition of FAK would augment and sustain the anti-proliferative effects of BMS-986158. Moreover, studies of 3D tumor spheroids in collagen demonstrated that the enhanced invasive potential of BET inhibitor DTP cells can be reversed by Defactinib.

### Combining FAK and BET inhibition prolongs survival and blocks metastatic colonization in vivo

To determine if the combination of BMS-986158 and Defactinib would be synergistic and tolerated in vivo we first evaluated subcutaneous xenografts. To mitigate toxicities that are often experienced by NSG mice, we used athymic nude mice for this study. Mice bearing A673 tumor xenografts were randomized to receive vehicle, BMS-986158, Defactinib, or the combination per the schema shown in **Figure 6A**. In contrast to earlier studies in NSG mice (**Fig 1**), single agent BMS-986158 did not inhibit tumor growth in nude mice (**Fig 6B**). Consistent with in vitro data, overall survival was superior when BMS-986158 was combined with Defactinib, as compared to vehicle (p=0.0001) or single agent BMS-986158 (p=0.0003), (**Fig 6B**) and no significant toxicity was observed with this drug combination (**Fig 6C**). However, the effect of Defactinib alone on subcutaneous tumor growth was not significantly augmented by addition of BMS-986158 indicating that, in nude mice, the combination of both drugs is no more effective at controlling bulky disease than Defactinib alone.

**Figure 6.**
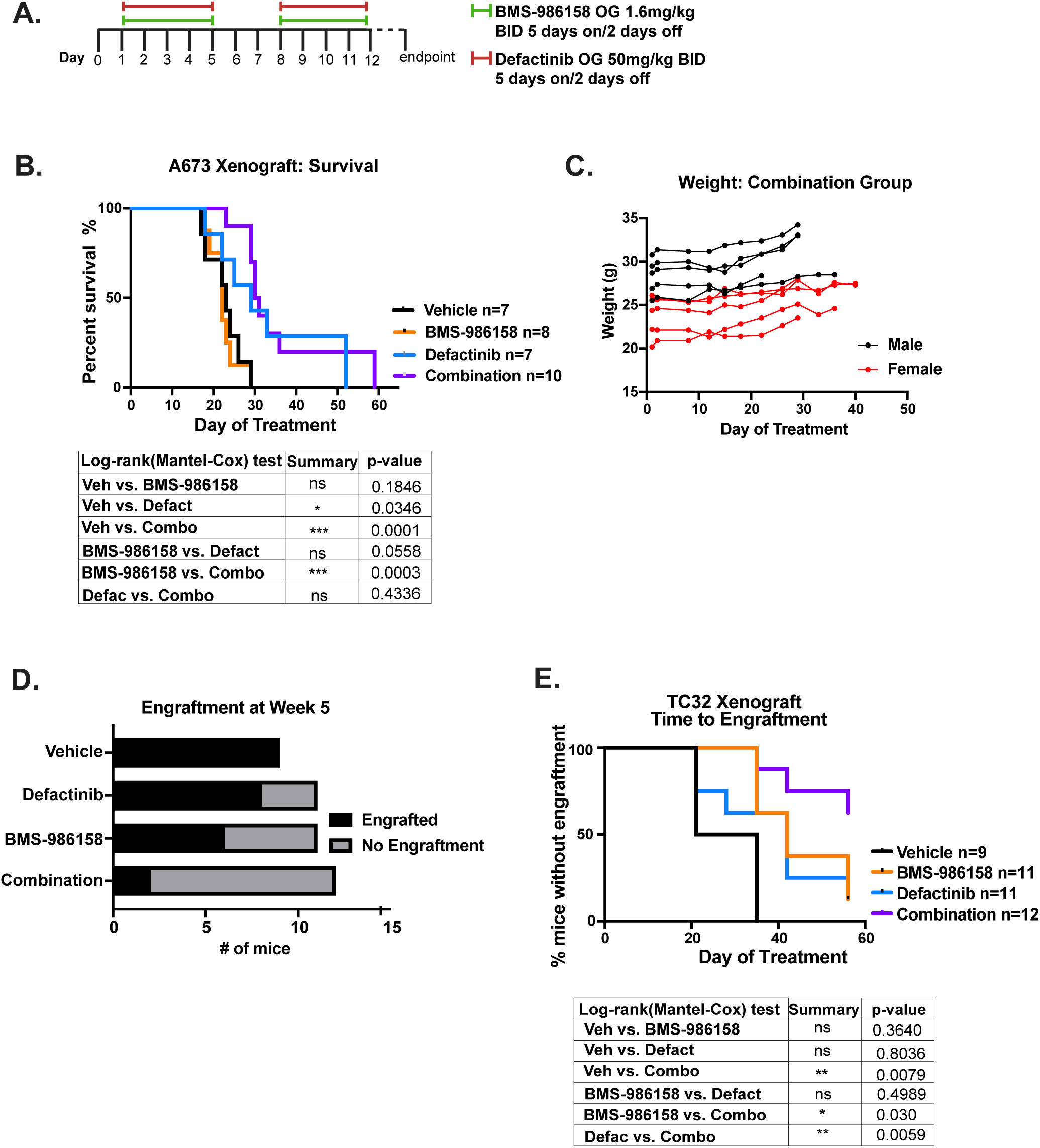
Combining FAK and BET inhibition prolongs survival and blocks metastatic colonization in vivo. **A.** Dosing schema of xenograft studies in athymic nude mice including dosing, route, and schedule of each drug. **B.** Kaplan-Meier overall survival curves as measured by Log-rank test. Table of p-values below. **C.** Weight of combination group over time. **D.** Number of mice in each treatment group with evidence of engraftment by IVIS imaging at start of Week 5. Statistical significance of engraftment between each comparison was determined by Fischer’s exact test. **E.** Time to engraftment plotted as Kaplan-Meier. Table of log-rank p-values below.

Tumor:ECM crosstalk and ECM remodeling are critical for successful engraftment and colonization of disseminated tumors cells(56). Given the importance of FAK to tumor:ECM signaling, and BET activity to cell proliferation, we next used a tail vein injection model to test whether the combination of FAK and BET inhibition would block colonization of microscopic tumor foci. We have observed that, unlike other EwS cell lines, TC32 cells can reproducibly form disseminated tumors in athymic nude mice when injected via tail vein. We thus injected 1x10^6^ TC32 cells into recipient mice and began drug treatment as per **Fig 6A** 72 hrs after cell inoculation. Treatment continued for up to 8 weeks and tumor colonization was monitored via serial bioluminescence. By the start of week 5, 100% of vehicle mice demonstrated tumor engraftment. At this same timepoint, we observed that 72.7% (8/11) of mice receiving single agent Defactinib and 54.5% (6/11) of mice receiving single agent BMS-986158 had engrafted. Significantly, at this same time point only 20% (2/12) of the combination group had evidence of engraftment (p=0.0002, Fischer’s exact test as compared to vehicle) (**Fig 6D**).

Colonization of disseminated EwS cells was also significantly inhibited by BMS-986158 (p=0.0379, Fisher’s exact test) as a single agent as compared to vehicle by week 5, confirming BMS-986158 activity in the setting of microscopic disease. While Defactinib delayed tumor colonization, it did not reach the threshold of statistical significance (p=.2184, Fisher’s exact test) (**Fig 6D**). Treatment was subsequently continued for an additional 4 weeks, and after 8 weeks of total treatment, the combination continued to significantly block tumor cell engraftment, as over 60% of mice did not show evidence of tumor colonization (**Fig 6E**). Again, we did not observe any significant toxicities in the mice receiving the combination of BMS-986158 and Defactinib. Thus, these studies confirm the safety and potential synergistic anti-tumor effects of BMS-986158 and Defactinib in combination for patients with recurrent EwS, especially in the setting of disseminated minimal residual disease.

## Discussion

In these studies, we have confirmed that the clinical grade BET inhibitor BMS-986158, which was the first BET inhibitor used in pediatric patients (NCT03936465), has anti-tumor efficacy in EwS. Like the tool compound JQ-1, it potently reverses the EWS::FLI1-activated transcriptional signature and this is accompanied, in the short term, by striking anti-proliferative and cytotoxic effects. In addition, our studies validate that combining BET inhibition with IGF1R/PI3K/AKT pathway inhibition leads to synergistic anti-proliferative effects, as previously reported(16). However, studies of this combination in xenograft models were found to be highly toxic, likely precluding their clinical use in combination in heavily pretreated patients. Importantly, our data show that under the pressure of BET inhibitor monotherapy, surviving EwS tumor cells undergo transcriptional rewiring, creating DTP cell populations that are vulnerable to FAK inhibition. Dual targeting of FAK and BET enhances cytotoxicity and blocks invasion in vitro and inhibits colonization of disseminated tumor cells in vivo. Our studies highlight that interrogating mechanisms of emergent tolerance to single agent epigenetic modifiers can inform biologically rational drug combinations that have the potential to augment and sustain anti-tumor efficacy.

DTP cell populations utilize complex adaptive mechanisms to survive and persist under therapeutic selective pressures. Specific mechanisms that have been elucidated include transcriptional and epigenetic reprogramming (19, 20, 42), metabolic alterations (57), transitions in cell state identity (18), and modification to the TME (21–23, 44). These alterations in DTP cell populations are typically reversible; however, investigating the mechanisms driving their development can reveal potential therapeutic vulnerabilities (20, 21, 23, 42). Transcriptional rewiring has been previously described as a primary mechanism of BET inhibitor escape. In AML, persistent BET inhibition led to loss of PRC2-mediated repression of transcriptional networks that restored *Myc* transcription and activated Wnt-signaling in the face of continued BRD4 inhibition (19). Further, in neuroblastoma, BET inhibition induced global enhancer rewiring leading to subsequent activation of multiple receptor tyrosine kinases, in particular PI3K (42). Mechanisms of BET inhibitor resistance in EwS have yet to be elucidated and transcriptional rewiring as a BET inhibitor escape mechanism has not been described in these tumors. Previous studies demonstrated that BRD4 inhibitor–resistant cell populations regained proliferative capacity following prolonged single-agent treatment. These resistant populations exhibited elevated CDK9 activity, facilitating BRD4-independent transcription (17). Our current studies now show that in response to prolonged BET inhibition, EwS cells undergo phenotypic and transcriptomic changes to a new cell state. In particular, DTP cells transcriptionally upregulate gene programs involved in tumor-matrix signaling that are normally subjected to EWS::FLI1-mediated repression, rendering cells susceptible to FAK inhibition.

Published studies have previously identified the importance of FAK signaling to EwS survival, proliferation, and invasion in vitro and in vivo (53, 58, 59). Importantly, there is increasing evidence that FAK signaling is a driver of DTP states in other solid tumors, including non-small cell lung cancer (NSCLC) (60) and melanoma (61), supporting our own finding that FAK is a vulnerability in BET inhibitor DTP cells in EwS. In melanoma, DTP populations that emerge following dual MEK and BRAF inhibition undergo cell state changes to a neural crest-like state, where FAK signaling is highly enriched (61). Importantly, and consistent with our own work, combining FAK inhibition with dual MEK and BRAF inhibition delayed engraftment of minimal residual disease populations and tumor formation in melanoma models (61). Together, these results suggest that altered FAK signaling may play a seminal role in cell state changes that mediate drug resistance, and that are induced in response to diverse targeted agents.

Activation of FAK signaling is a hallmark in many different cancer types (62). In addition, there is growing evidence that dysregulated FAK signaling in response to microenvironmental stress can contribute to tumor stroma fibrosis, leading to therapeutic resistance and tumor progression (62). The recent development of clinically relevant and therapeutic FAK inhibitors has resulted in multiple early phase clinical trials, and recent studies have shown the safety and efficacy of Defactinib as both a single agent, in combination with other agents (63), and in combination with radiotherapy (64). Importantly, Defactinib was recently granted FDA approval for use in combination with a dual MEK/RAF inhibitor for KRAS-mutated recurrent low-grade serous ovarian cancer (NCT06072781). Our work adds to the growing body of literature that supports targeting FAK in combination with other anti-cancer therapeutics as a means of augmenting anti-tumor efficacy.

While BET inhibitors and other epigenetic modifying agents are promising agents in EwS, the effects on the EWS::FLI1 transcriptional signature are complex. Initial studies of BET inhibition in EwS primarily focused on acute changes to the EWS::FLI1-activated transcriptional signature, where strong reversal was observed (12–15). However, changes to the repressed transcriptional signature, or how the EWS::FLI1 transcriptional signature evolves over time, have not been investigated. We found that despite restored proliferation, strong repression of the EWS::FLI1-activated signature was maintained in DTP populations. By contrast, we observed a striking difference in expression of the EWS::FLI1-repressed signature over time. While BET inhibition had little immediate impact on the repressed signature acutely, we observed that EWS::FLI1-repressed genes were strongly activated in DTP populations. This suggests that transcriptional rewiring of the BET inhibitor tolerant cells at least partially reversed the capacity of the fusion to repress its target genes. Many of these targets encode proteins that are critical for matrisomal remodeling, such as collagens and tenascin-C that bind and activate integrins on tumor cell membranes and initiate the FAK signaling cascade (55). Thus, selective reactivation of EWS::FLI1-repressed matrisomal genes likely contributes to the dependency of BET inhibitor DTP cells on FAK activity.

BET inhibitors and other epigenetic therapies remain promising treatments for EwS and a range of adult and pediatric solid tumors. However, drug tolerance is a significant barrier to clinical success, underscoring the need to study resistance mechanisms and identify rational combination strategies (65). Our findings emphasize the importance of investigating acquired drug-tolerant states, which often reveal novel vulnerabilities and therapeutic targets. By defining the shared features of these states, we can gain insights that can guide more effective therapeutic combinations. Based on the current work, we propose that dual BET and FAK inhibition may represent a rational approach to targeting minimal residual disease states in patients with disseminated EwS.

## Acknowledgements

The authors thank members of the Lawlor laboratory for insightful discussion, and the staff at the Seattle Children’s Research Institute Microscopy and Histopathology CoLab and Office of Animal Care. We would also like to thank the co-investigators of the Pediatric Stand Up to Cancer Catalyst Grant for their insightful discussions. This research was sponsored by a Pediatric Stand Up To Cancer Catalyst Grant supported by Bristol-Myers Squibb (Grant Number: SU2C#6143). Additional support for this work is gratefully acknowledged and was provided by the following sources: NIH/NCI 5K12CA076930 (SSG), NIH/NCI Loan Repayment Grant 1L40CA264716 (SSG), Alex’s Lemonade Stand Young Investigator Award (SSG), QuadW Foundation ASCO Conquer Cancer Young Investigator Award (SSG), Hyundai Hope on Wheels Young Investigator Award (SSG), CURE Childhood Cancer Foundation T2C Grant (ERL), Nick Teddy Foundation (ERL), Sam Day Foundation (ERL), and R01CA273081 (TSG).

---

**Supplementary Figure 1.**
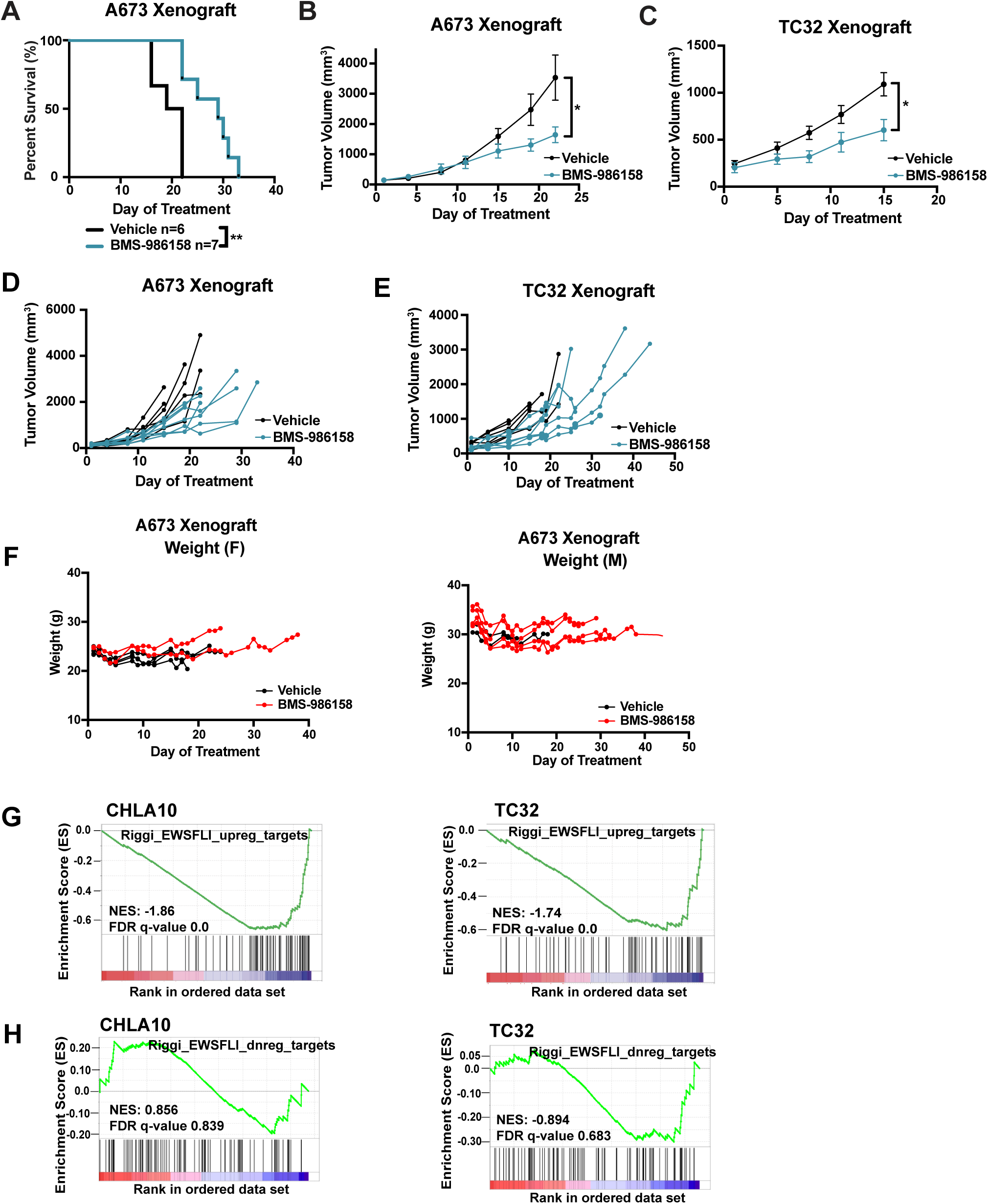
NSG xenograft models treated with BMS-986158 or vehicle. **A.** Kaplan-Meier overall survival curves of A673 xenografts treated with BMS-986158 or vehicle (p=0.0174). Average tumor volume of **B.** A673 (p=0.0333) and **C.** TC32 (p=0.0273) xenografts treated as measured by caliper 3x/week. Individual animal tumor volume plotted for **D.** A673 and **E.** TC32 xenografts. **F.** Weight of treatment and vehicle A673 xenografts. RNA-seq of CHLA10 and TC32 cells treated with 5nM BMS-986158 for 72hrs as compared to DMSO control. GSEA of 100 enhancer-driven genes **G.** activated or **H.** repressed by EWS::FLI1 (10).

**Supplementary Figure 2.**
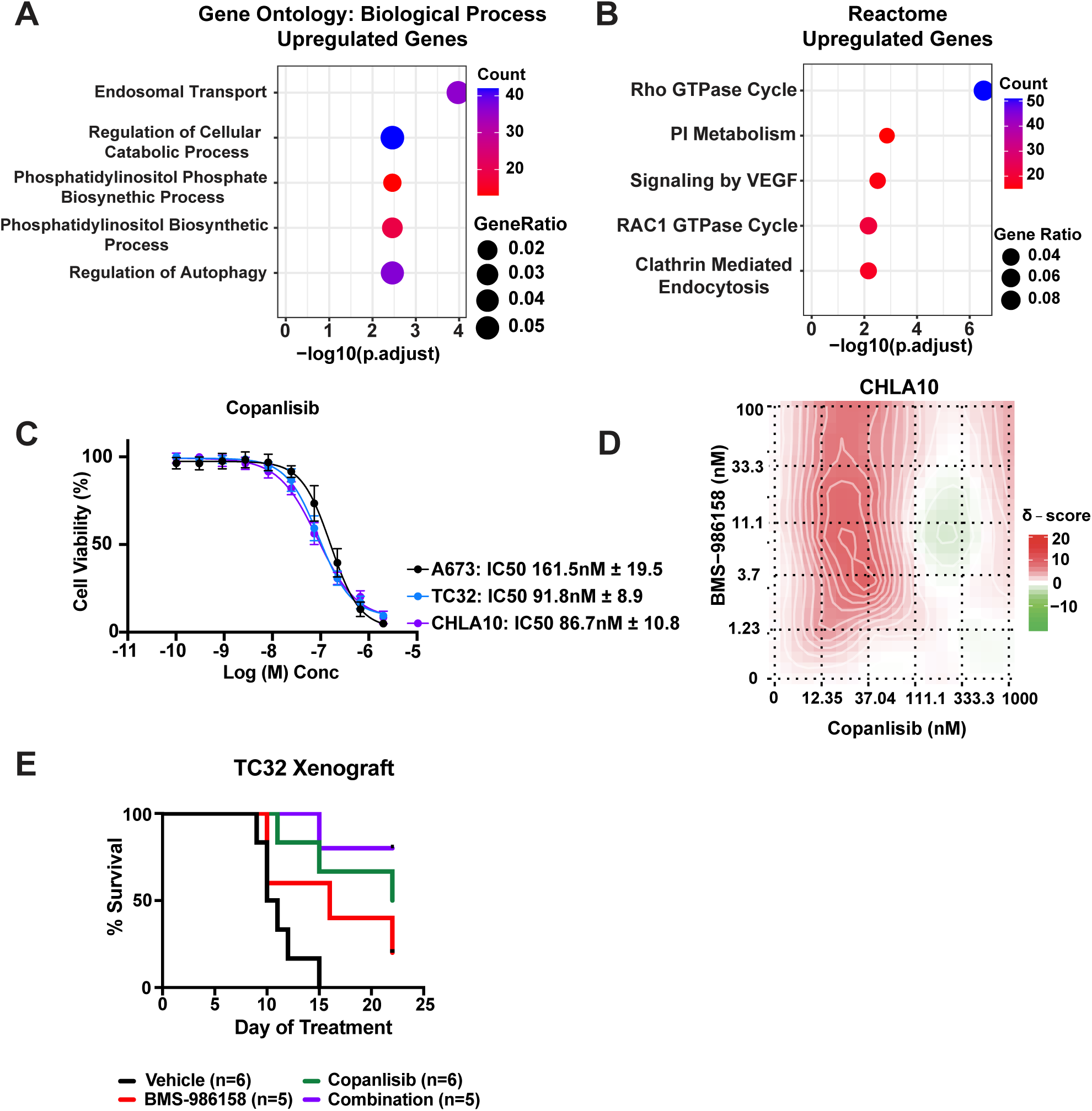
**A.** Reactome (Enrichr) and **B.** Gene Ontology Biological Process (Enrichr) over-representation analysis of the 889 genes that are upregulated in A673, CHLA10, and TC32 following 72hr treatment with BMS-986158. **C.** Dose response curves of multiple EwS cell lines treated with serial dilutions of Copanlisib from 0.11nM to 2uM, or relevant DMSO controls. **D.** Bliss Synergy (26) plots of BMS-986158 (0.41nM-100nM, serial dilutions 1:3) and Copanlisib (12.35nM-1uM, serial dilutions 1:3) exposed CHLA10 cells. **E.** Kaplan-Meier overall survival curves of TC32 xenografts treated with vehicle, BMS-986158, Copanlisib or combination, excluding mice in the combination group who were humanely euthanized at day 14 given weight loss >15%.

**Supplementary Figure 3.**
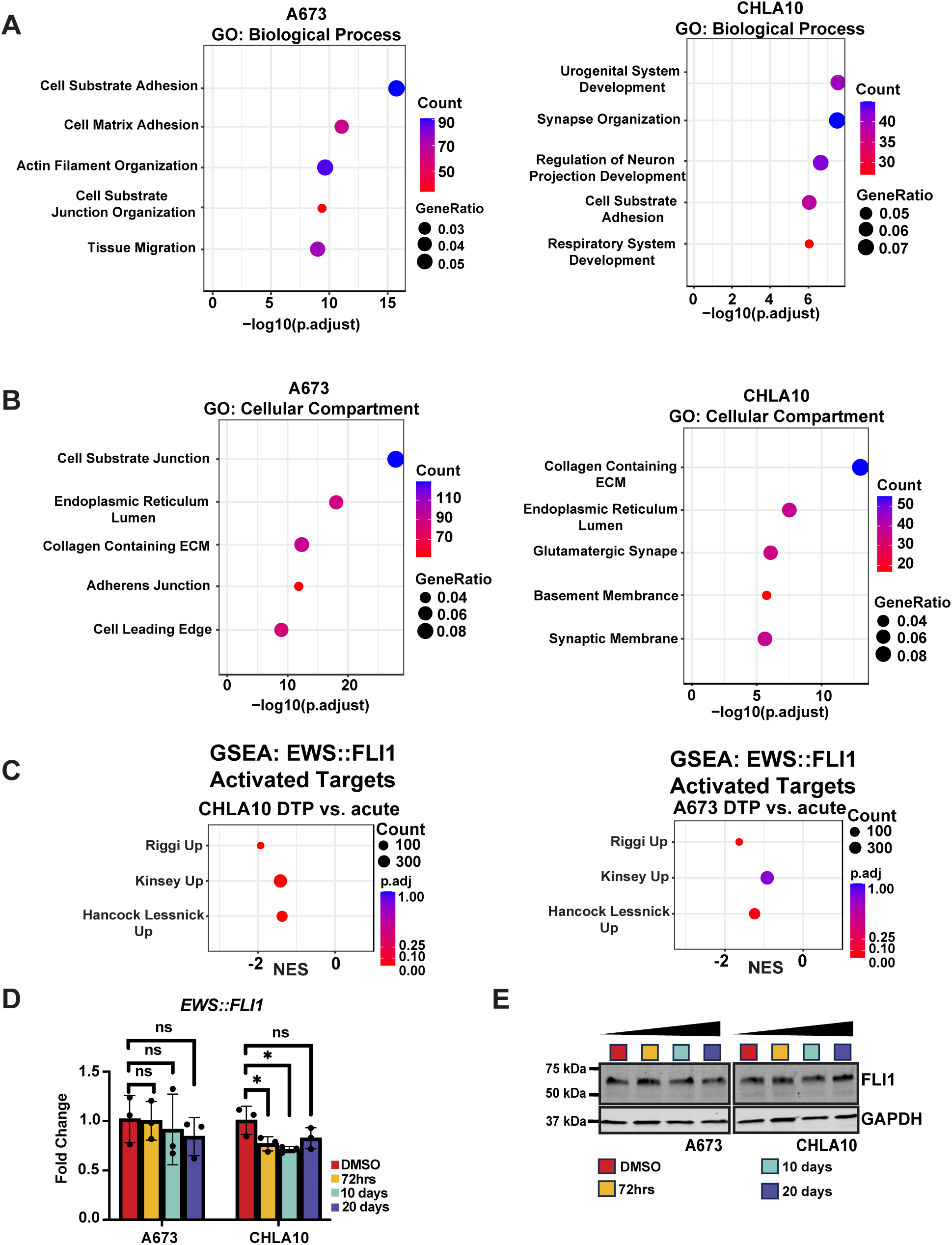
Differential expression by RNA-seq of cell lines treated with 5nM BMS-986158 for 20 days as compared to 5nM BMS-986158 for 72hrs was performed in A673 and CHLA10 cells. **A.** Gene Ontology, Biological Processes (Enrichr) and **B.** Gene Ontology, Cellular Components (Enrichr) over-representation analysis of the 1801 genes in A673 cells and 694 genes in CHLA10 cells that are significantly upregulated with 20 days vs. 3 days of BMS-986158 treatment. **C.** GSEA of activated EWS::FLI1 signatures (10, 66, 67). **D.** *EWS::FLI1* gene expression (n=3 with three technical replicates per experiment) by RT-qPCR and **E.** immunoblot of EWS::FLI1 protein expression (representative image of n=2) in A673 and CHLA10 cells treated with 72hrs, 10 days, or 20 days of 5nM BMS-986158.

**Supplementary Figure 4.**
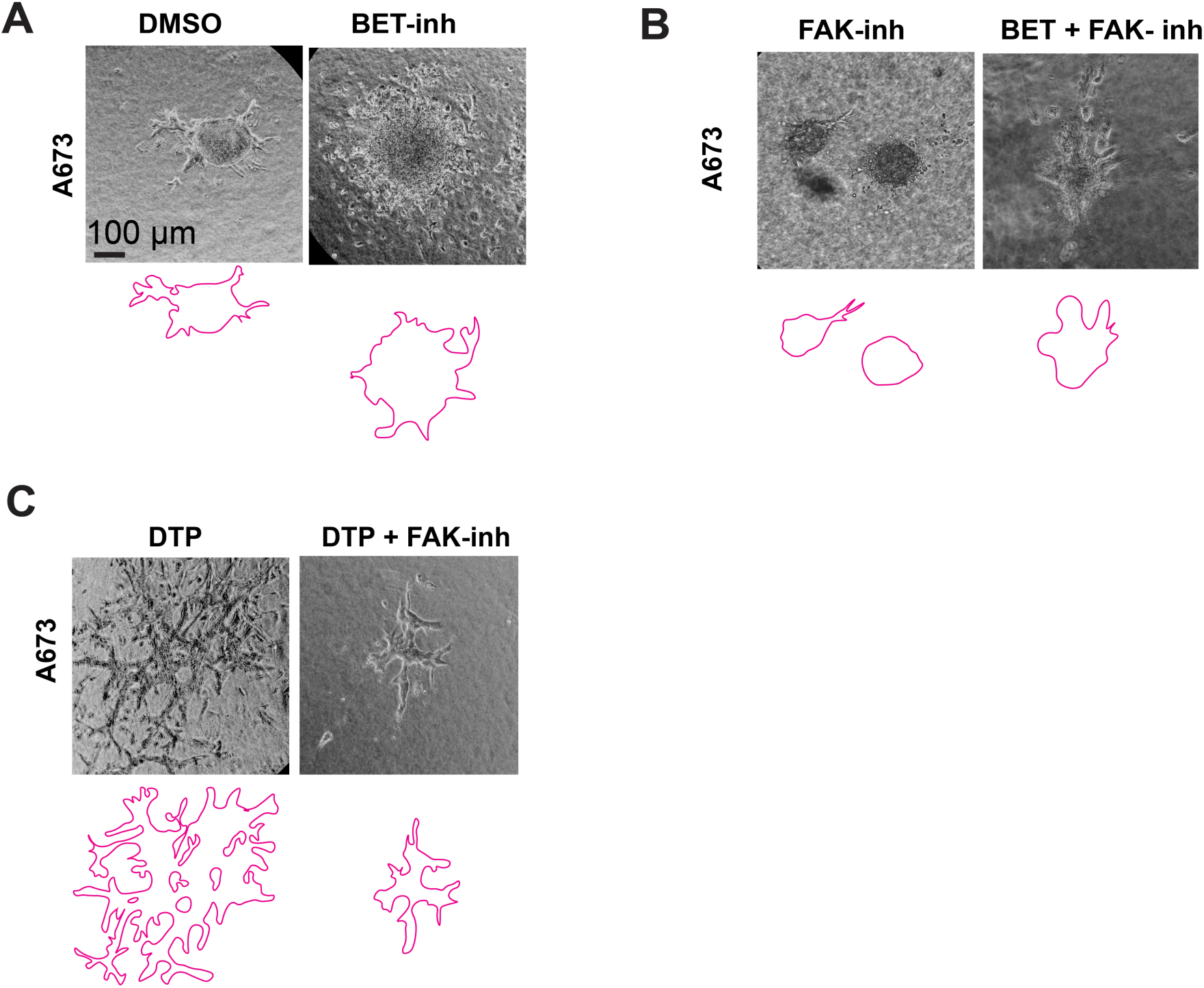
A673 naïve spheroids were embedded in rat tail collagen matrix. Phase contrast imaging was performed 6 days later to assess invasion through a collagen matrix. Embedded spheroids were treated with **A.** 5nM BMS-986158 vs. DMSO or **B.** 5uM Defactinib vs. the combination of 5nM BMS-986158 and 5uM Defactinib **C.** A673 DTP spheroids were embedded in rat tail collagen with or without 5uM Defactinib.

